# An integrated anatomical, functional and evolutionary view of the *Drosophila* olfactory system

**DOI:** 10.1101/2025.01.16.632927

**Authors:** Richard Benton, Jérôme Mermet, Andre Jang, Keita Endo, Steeve Cruchet, Karen Menuz

## Abstract

The *Drosophila melanogaster* olfactory system is one of the most intensively studied parts of the nervous system in any animal. Composed of ∼60 independent olfactory neuron classes, with several associated hygrosensory and thermosensory pathways, it has been subject to diverse types of experimental analyses. However, synthesizing the available data is limited by the incompleteness and inconsistent nomenclature found in the literature. In this work, we first “complete” the peripheral sensory map through the identification of a previously uncharacterized antennal sensory neuron population expressing Or46aB, and the definition of an exceptional “hybrid” olfactory neuron class comprising functional Or and Ir receptors. Second, we survey developmental, anatomical, connectomic, functional and evolutionary studies to generate an integrated dataset of these sensory neuron pathways – and associated visualizations – creating an unprecedented comprehensive resource. Third, we illustrate the utility of the dataset to reveal relationships between different organizational properties of this sensory system, and the new questions these stimulate. These examples emphasize the power of this resource to promote further understanding of the construction, function and evolution of these neural circuits.

## Introduction

Sensory regions of the nervous system are, by virtue of their peripheral location and molecularly-distinct cell types, particularly amenable for developmental, anatomical and physiological investigations to obtain a holistic view of the construction and function of neural circuits. Amongst model sensory systems, the olfactory pathways of *Drosophila melanogaster* are some of the most intensively studied (Benton, 2022; Grabe and Sachse, 2018; Jefferis and Hummel, 2006; Su et al., 2009; Vosshall and Stocker, 2007) (Figure 1A).

**Figure 1.**
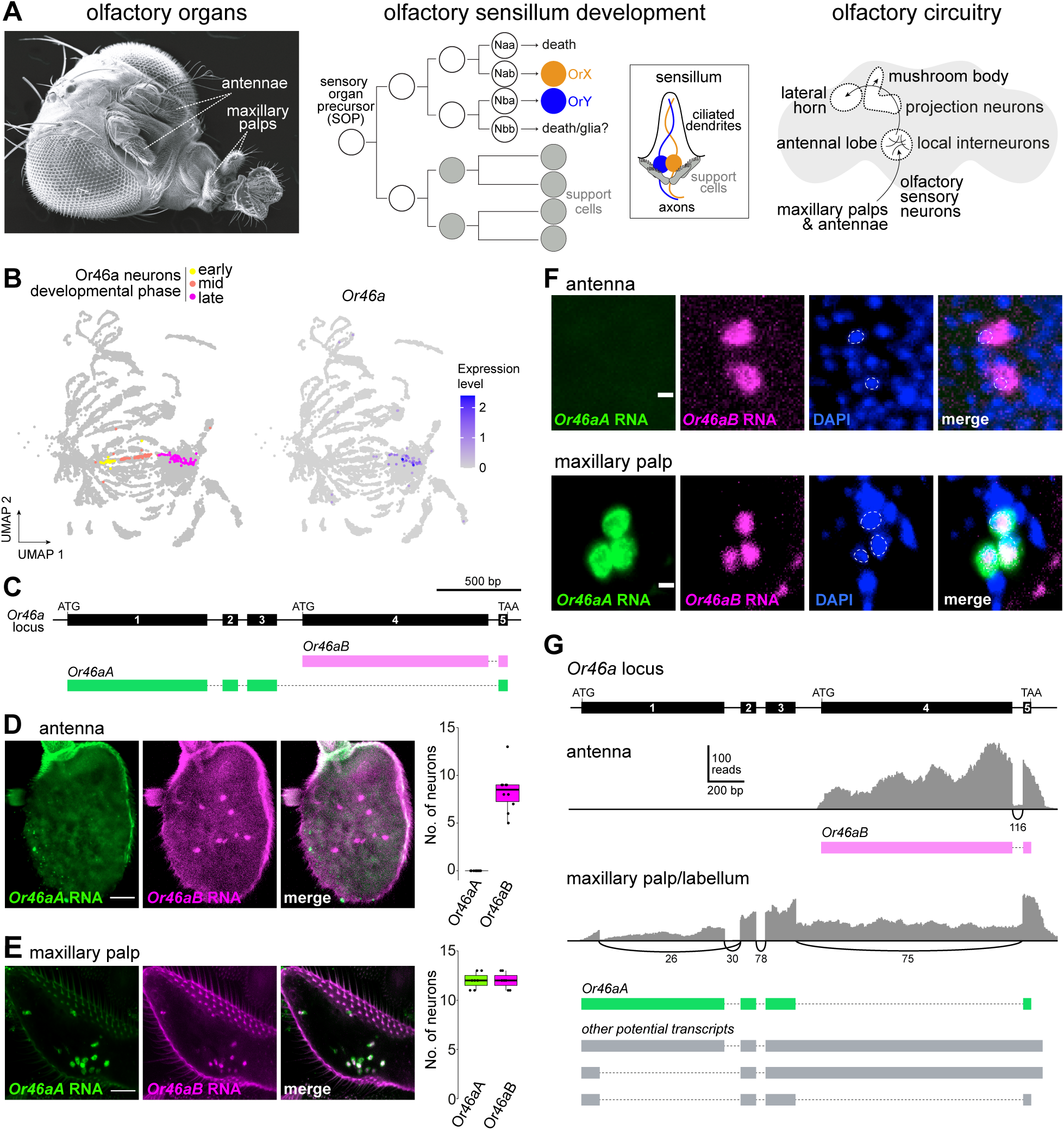
A new antennal olfactory sensory neuron population. (A) Schematic of *D. melanogaster* olfactory system anatomy, development and circuitry (see text for details). The scanning electron micrograph (left) was adapted from (Benton and Dahanukar, 2011) (copyright © Cold Spring Harbor Laboratory Press). (B) UMAP of an snRNA-seq atlas of developing antennal neurons colored for developmental phase of the Or46a neurons (“early” = 18-30 h after puparium formation (APF)), “mid” = 36-48 h APF, “late” = 56-80 h APF) (left) and expression of *Or46a* transcripts (right). Data from (Mermet et al., 2025). Gene expression levels, here and in other UMAPs, are residuals from a regularized negative binomial regression and have arbitrary units. (C) Structure of the *Or46a* locus and the transcript isoforms for *Or46aA* and *Or46aB*. (D-E) RNA FISH with isoform-specific probes for *Or46aA* and *Or46aB* in a whole-mount antenna (D) and maxillary palp (E). Scale bars, 25 μm. Quantifications of neuron numbers are shown on the right. Box plots show median (thick line), first and third quartiles, while whiskers indicate data distribution limits, overlaid with individual data points (*n* = 10 (D) and 7 (E)). (F) High-magnification images of RNA FISH for *Or46aA* and *Or46aB* in an antenna and a maxillary palp. Dashed lines outline the nuclei (stained with DAPI), revealing greater nuclear sequestration of *Or46aB* in the maxillary palp neurons, compared to *Or46aA* transcripts, or to *Or46aB* transcripts in the antenna. Scale bars, 3 μm. (G) *Or46a* isoform expression analyzed from bulk RNA-seq data of antennal and maxillary palp/labellar tissue (Bontonou et al., 2024). Top: structure of the *Or46a* locus. Sashimi plots generated with IGV (Thorvaldsdottir et al., 2013) showing mapped reads (grey) from the indicated tissue transcriptomes aligned to the *Or46a* locus. Quantification of splice junction mapping reads are indicated beneath the plots, and the predicted transcript isoforms in each tissue are shown below. Potential transcripts in the palp shown in grey are unlikely to encode functional receptor proteins (see Results).

Odor-sensing occurs in two bilaterally-symmetric pairs of peripheral organs, the maxillary palps and antennae. These appendages are covered with hundreds of porous sensory hairs, or sensilla, of distinct morphologies (basiconic, trichoid, intermediate, coeloconic) (Nava Gonzales et al., 2021; Shanbhag et al., 1999, 2000; Shanbhag et al., 1995). Sensilla house the ciliated dendrites of 1-4 olfactory sensory neurons (OSNs), each of which expresses a specific type of odor-binding sensory receptor (or occasionally receptors) that recognize a defined set of volatile chemicals (Couto et al., 2005; de Bruyne et al., 1999; de Bruyne et al., 2001; Fishilevich and Vosshall, 2005; Munch and Galizia, 2016; Silbering et al., 2011). Approximately 25 functional classes of olfactory sensilla on the antenna and maxillary palp can be identified by the stereotypical receptor expression patterns and odor response profiles of the neurons they house (Couto et al., 2005; de Bruyne et al., 1999; de Bruyne et al., 2001; Grabe et al., 2016; van der Goes van Naters and Carlson, 2007; Yao et al., 2005).

Olfactory receptors belong to two families of ligand-gated ion channels: the Odorant receptors (Ors), the founder members of the seven transmembrane domain ion channel (7TMIC) superfamily (Benton and Himmel, 2023; Butterwick et al., 2018; Clyne et al., 1999b; Del Marmol et al., 2021; Gao and Chess, 1999; Himmel et al., 2023; Sato et al., 2008; Vosshall et al., 1999; Wicher et al., 2008), and the Ionotropic receptors (Irs), which are distantly-related to ionotropic glutamate receptors (iGluRs) (Benton et al., 2009). Both Ors and Irs function in known (or presumed) heterotetrameric complexes composed of “tuning” receptor subunits that are thought to directly bind odors, and subunits of one or more broadly-expressed co-receptors (Orco for Ors (Larsson et al., 2004); Ir8a, Ir25a and Ir76b for Irs (Abuin et al., 2011; Vulpe and Menuz, 2021)). Other tuning Ir subunits form hygrosensory and thermosensory receptors with Ir25a and Ir93a co-receptors expressed by sensillar neurons within specialized antennal structures: the sacculus, a three-chambered internal pocket that also houses some olfactory neurons (Ai et al., 2010; Vulpe et al., 2021)), and the arista, an elongated cuticular projection (Budelli et al., 2019; Enjin et al., 2016; Frank et al., 2017; Gallio et al., 2011; Knecht et al., 2017; Knecht et al., 2016; Marin et al., 2020). Finally, a few “Gustatory receptors” (Grs), which are also 7TMICs, function in antennal neurons in CO_2_ detection (Jones et al., 2007; Kwon et al., 2007) and thermosensation (Ni et al., 2013).

During development, each sensillum derives from an individual sensory organ precursor (SOP) cell in the pupal antennal imaginal disk, which undergoes three stereotyped rounds of division to produce four support cells and four sensory neuron precursors termed Naa, Nab, Nba and Nbb (Chai et al., 2019; Endo et al., 2007; Endo et al., 2011; Jefferis and Hummel, 2006; Rodrigues and Hummel, 2008) (Figure 1A). (In many coeloconic lineages the Nbb precursor is thought to differentiate as a glial cell (Endo et al., 2007; Rodrigues and Hummel, 2008; Sen et al., 2005).) Support cells have diverse roles in synthesizing and shaping the sensillar cuticle during development (Ando et al., 2019; Schmidt and Benton, 2020), as well as secreting perireceptor proteins into the sensillar lymph that bathes neuronal dendrites, where they can contribute to sensory responses (Larter et al., 2016; Sun et al., 2018; Xu et al., 2005). Sensory neuron precursors are thought to express unique combinations of transcription factors that, together with asymmetric Notch activity between daughter cells of each division, result in the unique terminal identities of the olfactory neurons (Barish and Volkan, 2015; Chai et al., 2019; Endo et al., 2007; Endo et al., 2011; Mermet et al., 2025). In most sensillar classes, one or more sensory neuron precursors stereotypically undergo programmed cell death, leaving fewer than four functional neurons in mature sensilla (Endo et al., 2007; Endo et al., 2011; Prieto-Godino et al., 2020; Sen et al., 2004).

Populations of sensory neurons expressing the same receptor(s) innervate a specific glomerulus in the antennal lobe, the initial processing center in the brain (Couto et al., 2005; Fishilevich and Vosshall, 2005; Gao et al., 2000; Silbering et al., 2011; Vosshall et al., 2000). Here these sensory neurons synapse with local neurons (LNs), which mediate interglomerular interactions (Chou et al., 2010; Wilson, 2013) and projection neurons (PNs), which transmit sensory information to higher processing centers, the mushroom body and lateral horn (Bates et al., 2020; Marin et al., 2020; Marin et al., 2002; Schlegel et al., 2021; Wong et al., 2002) (Figure 1A).

The global view of the organization and function of the *D. melanogaster* olfactory system has emerged from diverse experimental approaches over the past 25 years. Odor response profiles of nearly all receptors and/or sensory neurons have been obtained through measurement of odor-evoked activity *in vivo* by extracellular electrophysiological recordings from individual sensilla (e.g., (de Bruyne et al., 1999; de Bruyne et al., 2001; Hallem and Carlson, 2006; Yao et al., 2005)), optical imaging of activity in sensory neuron axonal termini in glomeruli (e.g., (Silbering et al., 2011; Wang et al., 2003)) and/or through characterization of receptors in heterologous expression systems (e.g., (Ruel et al., 2021; Sato et al., 2008)). *In situ* analysis of the expression of endogenous receptors or transgenic promoter reporters (Benton et al., 2009; Couto et al., 2005; Fishilevich and Vosshall, 2005; Grabe et al., 2016; Silbering et al., 2011) has been complemented with comprehensive, high resolution transcriptomic analyses of OSNs and PNs (Arguello et al., 2021; Li et al., 2017; Li et al., 2020; McLaughlin et al., 2021). Receptor promoter transgenic reporters have also enabled neuronal tracing to produce a near-complete, neuron-to-glomerulus map (Couto et al., 2005; Fishilevich and Vosshall, 2005; Silbering et al., 2011), which has recently been greatly extended by electron microscopic (EM) analyses that also offer insights into the glomerular microcircuitry of sensory neurons, LNs and PNs (Bates et al., 2020; Marin et al., 2020; Rybak et al., 2016; Schlegel et al., 2021; Tobin et al., 2017), as well as the innervations of PNs in higher brain regions (Bates et al., 2020; Jefferis et al., 2007; Marin et al., 2020; Schlegel et al., 2021). Insights into how this circuitry forms have been discovered through a wealth of forward and reverse molecular genetic investigations of OSN and PN development (Barish and Volkan, 2015; Brochtrup and Hummel, 2011; Hong and Luo, 2014; Jefferis and Hummel, 2006). The behavioral role(s) of many individual sensory pathways have been revealed by genetic manipulations of receptors, as well as artificial inhibition or activation of the neurons in which they are expressed (e.g., (Ai et al., 2010; Stensmyr et al., 2012; Suh et al., 2004; Tumkaya et al., 2022; Wu et al., 2022)). Finally, comparative analysis of the *D. melanogaster* olfactory system with that of other drosophilids and more distantly-related insect species has begun to uncover how individual sensory pathways diverge structurally and/or functionally during evolution (Auer et al., 2020; Dekker et al., 2006; Depetris-Chauvin et al., 2023; Hansson and Stensmyr, 2011; Prieto-Godino et al., 2016; Prieto-Godino et al., 2017; Ramdya and Benton, 2010; Takagi et al., 2024; Zhao and McBride, 2020).

These numerous investigations on *D. melanogaster*’s olfactory pathways provide essential resources for the field. However, integration of information across different studies can be difficult due to conflicting assignment of some receptors to neuron types and sensillar classes, inconsistent naming of antennal lobe glomeruli, and ongoing updates to the olfactory map. In this work, we first “complete” this map through the discovery of a previously undescribed antennal OSN type, which resolves long-known inconsistencies in sensillar identification. We also reveal a neuron that relies on both Ir and Or tuning receptors, the only such “hybrid” olfactory neuron known in *D. melanogaster*. These findings spurred us to compile an integrated data resource to overcome the dispersal of pertinent information with disparate anatomical and molecular naming across the literature. We also created updated representations of both the complete sensillar classes and the antennal lobe glomeruli to serve as standardized references for the field.

## Results and Discussion

### A novel antennal Or sensory neuron type

Within a single-nuclear RNA-sequencing (snRNA-seq) atlas of the developing antenna (Mermet et al., 2025), we observed a cell cluster expressing *Or46a* (Figure 1B). Transcripts for this gene had previously been observed by RT-PCR and in bulk RNA-seq datasets of the antenna (Clyne et al., 1999a; Menuz et al., 2014), but never assigned to a specific cell type. The *Or46a* locus encodes two receptors, Or46aA and Or46aB, which share the same C-terminus encoded by a common last exon (Figure 1C). Through RNA fluorescence *in situ* hybridization (FISH) with isoform-specific probes, we detected expression of transcripts for *Or46aB* in ∼8 neurons in the antenna, but not *Or46aA* (Figure 1D). As a control, we performed RNA FISH on maxillary palps, verifying that both *Or46a* probes detect the same neurons in this organ, as described previously (Ray et al., 2007) (Figure 1E). However, we observed that the signals of the two probes were spatially distinct (Figure 1F): *Or46aA* was detected both in the cytoplasm and the nucleus, while *Or46aB* appeared predominantly nuclear in palp OSNs (Figure 1F), despite being readily detected in the cytoplasm of antennal OSNs. This phenomenon is reminiscent of the nuclear retention of transcripts of downstream genes in tandem clusters of *Or*s in ants (Brahma et al., 2023).

To understand the reason for this differential location, we assessed transcripts arising from the *Or46a* locus in antenna and maxillary palp/labellum bulk transcriptomes (Bontonou et al., 2024) (Figure 1G). In the antenna, we detected transcripts only for *Or46aB*, as expected. In the maxillary palp/labellum transcriptome, we detected several alternative splicing events; many of these correspond to splicing events in *Or46aA*, as previously characterized by RT-PCR of full-length transcripts (Ray et al., 2007). Importantly, although we found transcripts including *Or46aB* exons we did not find any evidence for proper splicing between exons 4 and 5. This lack of splicing means that all transcripts with *Or46aB* exons contain a frameshift that renders exon 5 unable to encode for the essential ion channel pore region. We also observed sequences with an unusual alternative splicing event in the first exon of *Or46aA* that would prevent them encoding a functional receptor (Figure 1G). We suggest that many or all of these transcripts are aberrant splice variants initiating from the *Or46aA* promoter and likely fail to be exported efficiently from the nucleus or are rapidly degraded in the cytoplasm. The simplest interpretation of these data is that antennal neurons only express Or46aB protein, while maxillary palp neurons predominantly or only express Or46aA.

### “Completing” the olfactory map in the antenna and antennal lobe

We next sought the antennal sensillum class in which the newly-identified Or46aB neurons are housed, taking advantage of odor-to-neuron-to-sensillum maps defined by electrophysiological and histological analyses (Couto et al., 2005; de Bruyne et al., 2001; Grabe et al., 2016) and knowledge that Or46aB responds to methylphenols when expressed in heterologous neurons (Ray et al., 2014). We predicted that Or46aB is expressed in the antennal basiconic 6 (ab6) sensillar class “B” neuron (i.e., with the smaller spike amplitude) as this ab6B neuron responds strongly and selectively to methylphenols (de Bruyne et al., 2001; Hallem et al., 2004). The molecular identity of the ab6A neuron (i.e., with the larger spike amplitude) has been inconsistently described in the literature (see “Terminology” section in the Methods), but the best evidence is that this neuron class expresses Or13a, due to the similar odor-tuning profiles of ab6A neurons measured by single-sensillum recordings (de Bruyne et al., 2001) and Or13a neurons measured by calcium imaging (Galizia et al., 2010).

We tested this prediction through two-color RNA FISH using probes against these receptors, observing precise pairing of Or46aB and Or13a neurons (Figure 2A). We further investigated the neuronal composition and function of this sensillum through targeted electrophysiological recordings of sensilla labelled with GFP driven by *Or13a-Gal4*. Observation of basal spiking patterns confirmed the presence of two neurons, based upon their distinct spike amplitudes (Figure 2B), countering a previous claim that these sensilla house a single neuron (Lin and Potter, 2015). Profiling of the odor-evoked responses confirmed that the A neuron responds most strongly to *1*-octen-*3*-ol and robustly to *1*-hexanol, *E2*-hexenal, pentyl acetate, and *2*-heptanone, matching the profile of ab6A neurons previously defined by electrophysiological recordings (de Bruyne et al., 2001) and of Or13a neurons measured with calcium imaging (Galizia et al., 2010). As previously described for ab6B neurons (de Bruyne et al., 2001; Hallem et al., 2004), the neuron paired with Or13a neurons responds to methylphenols (Figure 2B-C), matching the response profile of heterologously-expressed Or46aB (Ray et al., 2014). Together these data support the proposal that Or13a and Or46aB are expressed in the originally-defined ab6 sensillum class (de Bruyne et al., 2001).

**Figure 2.**
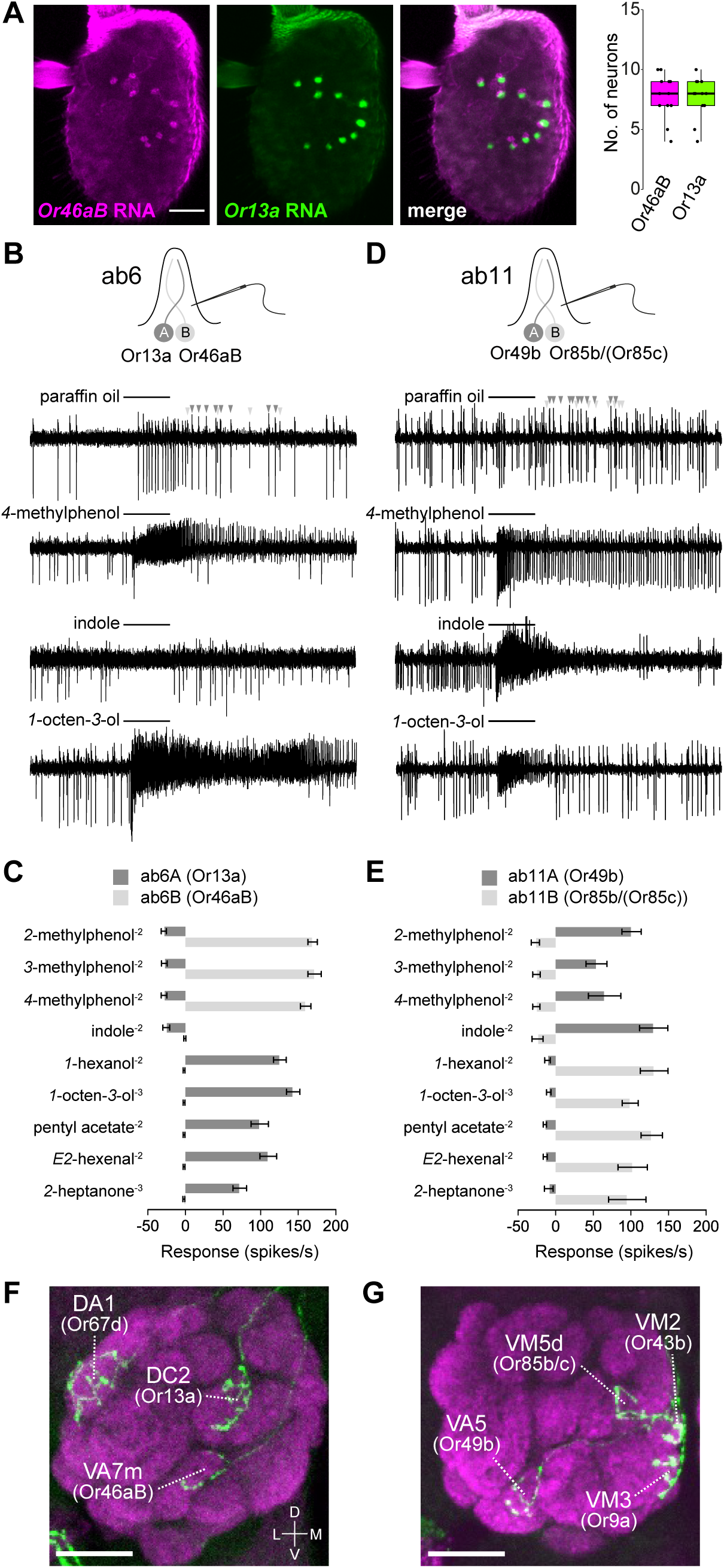
Molecular, functional and anatomical validation of ab6 and ab11 sensilla. (A) RNA FISH on a whole-mount antenna illustrating the pairing of Or46aB and Or13a neurons. Quantification of neuron numbers are shown on the right ( = 12). Scale bar, 25 μm. (B) Representative traces of single-sensillum recordings of GFP+ ab6 sensilla from *Or13a>mCD8:GFP* flies illustrating neuronal responses to the indicated odors (0.5 s stimulation time, black bars). In the top trace, two spike amplitudes, reflecting distinct neurons, are highlighted with dark and light grey arrowheads. (C) Quantification of odor-evoked responses in A (large spiking) and B (small spiking) neurons from ab6 sensilla. Odor dilutions (v/v in paraffin oil) are shown in superscript. Solvent-corrected responses (mean ± SEM) are shown. See Data S1 for spike counts and sample sizes. (D-E) As in (B-C), but for recordings of GFP+ ab11 sensilla from *Or49b>mCD8: GFP* flies. (F) Antennal lobe projections of clonally-marked OSNs visualized with GFP immunofluorescence (green) together with nc82 neuropil stain (magenta) revealing co-labeling of neurons innervating DC2 (Or13a) and VA7m (inferred to be Or46aB) glomeruli. Data were re-processed from (Endo et al., 2007); of 12 brains with DC2-labelled neurons, all had VA7m-labeled neurons (1 with weak labelling), strongly supporting the innervation patterns of the paired neurons in ab6. In this image, DA1 (Or67d) OSNs are also labeled, representing an independent clone in the at1 lineage. Scale bar, 20 μm. (G) Antennal lobe projections of clonally-marked OSNs innervating VA5 (Or49b) and VM5d (Or85b/(Or85c)) glomeruli. Data were re-processed from (Endo et al., 2007); of 4 brains with VA5-labelled neurons, 3 also had VM5d-labeled neurons, supporting the pairing of these neurons in ab11. In this image, VM2 (Or43b) and VM3 (Or9a) OSNs are also labeled, representing an independent clone in the ab8 lineage. Scale bar, 20 μm.

One complication with this assignment is that ab6B has previously been posited to express Or49b (e.g. (Couto et al., 2005; Grabe et al., 2016; Hallem et al., 2004)), likely because this receptor also responds to methylphenols (Hallem et al., 2004). Although it is possible that Or49b and Or46aB are co-expressed in ab6B, there is no evidence for this in our snRNA-seq datasets (Mermet et al., 2025). Moreover, we recently demonstrated using RNA FISH that Or49b neurons are paired with those expressing Or85b/(Or85c) (in this study, we place receptors in parentheses if their function is unclear) (Takagi et al., 2024). The simplest interpretation is that there are two discrete classes of sensilla, one with Or13a and Or46aB neurons and the other with Or85b/(Or85c) and Or49b neurons. These classes may have been conflated previously due to common sensitivity of both Or46aB and Or49b to methylphenols.

To validate that Or49b and Or85b/(Or85c) define a unique sensillum class, we used *Or49b-Gal4* to mark these sensilla with GFP and performed electrophysiological recordings with the same set of odors as above (Figure 2D-E). As expected, we found that the response profile of sensilla housing Or49b and Or85b/(Or85c) neurons is similar to those containing Or13a and Or46aB neurons. However, two key features indicate that the sensilla are distinct. First, methylphenols activate the A neuron in the Or49b sensilla (Figure 2D-E), but the B neuron in Or13a sensilla (Figure 2B-C), while odors such as *2-*heptanone and *1*-octen-*3*-ol activate the B neuron in Or49b sensilla, but the A neuron in Or13a sensilla. Second, the responses of Or13a and Or49b sensilla to indole, an odor reported to strongly activate Or49b (Ruel et al., 2021) differ: the A neuron in Or49b sensilla responds robustly to this odor, whereas neurons in Or13a sensilla do not (Figure 2B-E), as originally reported in ab6 (de Bruyne et al., 2001). Together, these data confirm that these receptors are expressed in two separate classes of sensilla, and that the ab6 sensilla response profile is matched best by the sensillum housing Or13a and Or46aB neurons. We propose to name the sensillum housing Or49b and Or85b/(Or85c) neurons ab11 (see the “Terminology” section in the Methods).

We next sought where Or46aB antennal OSNs project in the brain. Functional transgenic drivers for the Or46aB neuron have been difficult to generate (Couto et al., 2005; Tirian and Dickson, 2017), likely reflecting the unusual genomic organization of this locus (Figure 1C). This unfortunately prevents direct visualization of their glomerular target in the antennal lobe. However, we hypothesized that these neurons innervate the VA7m glomerulus. Three pieces of evidence support this possibility: VA7m is the last “orphan” glomerulus in the antennal lobe (Schlegel et al., 2021), i.e., without molecularly-defined sensory innervations. Second, the glomerulus is adjacent to the VA7l glomerulus, which is innervated by maxillary palp Or46aA neurons (Couto et al., 2005). Such an assignment aligns with evidence that evolutionarily closely-related receptors tend to be expressed in neurons that project to nearby glomeruli (Couto et al., 2005; Silbering et al., 2011). Most compellingly, clonal labelling of OSNs demonstrated that the sister neuron of Or13a – i.e., arising from the same SOP lineage, which we have now established is the Or46aB neuron (Figure 2A) – innervates VA7m (Figure 2F) (Endo et al., 2007). This neuron-to-glomerulus assignment effectively completes the antennal lobe map. Additionally, while reviewing data from (Endo et al., 2007), we found several examples of brains in which VA5 (Or49b) neurons are co-labeled with VM5d (Or85b/(Or85c)) neurons, supporting the pairing of these neurons in ab11 (Figure 2G). This co-labeling was previously over-looked as VM5d (Or85b/(Or85c)) neurons were mostly co-labeled with DM2 (Or22a/(Or22b)) neurons, corresponding to the co-housing of these OSN types in ab3.

### A “hybrid” olfactory pathway expressing a functional Or and Ir tuning receptor

Our snRNA-seq atlas (Mermet et al., 2025) revealed a second, previously-unreported expression pattern (Benton et al., 2009): weak expression of *Ir76a* in *Or35a*-expressing cells that correspond to the B neurons in antennal coeloconic 3 (ac3) sensilla (Figure 3A). (Stronger *Ir76a* expression was detected in the ac4 Ir76a neuron (Benton et al., 2009; Mermet et al., 2025)). We confirmed these transcriptomic data *in vivo* using RNA FISH, which detected *Ir76a* transcripts in several, though not all, *Or35a* ac3B neurons (Figure 3B).

**Figure 3.**
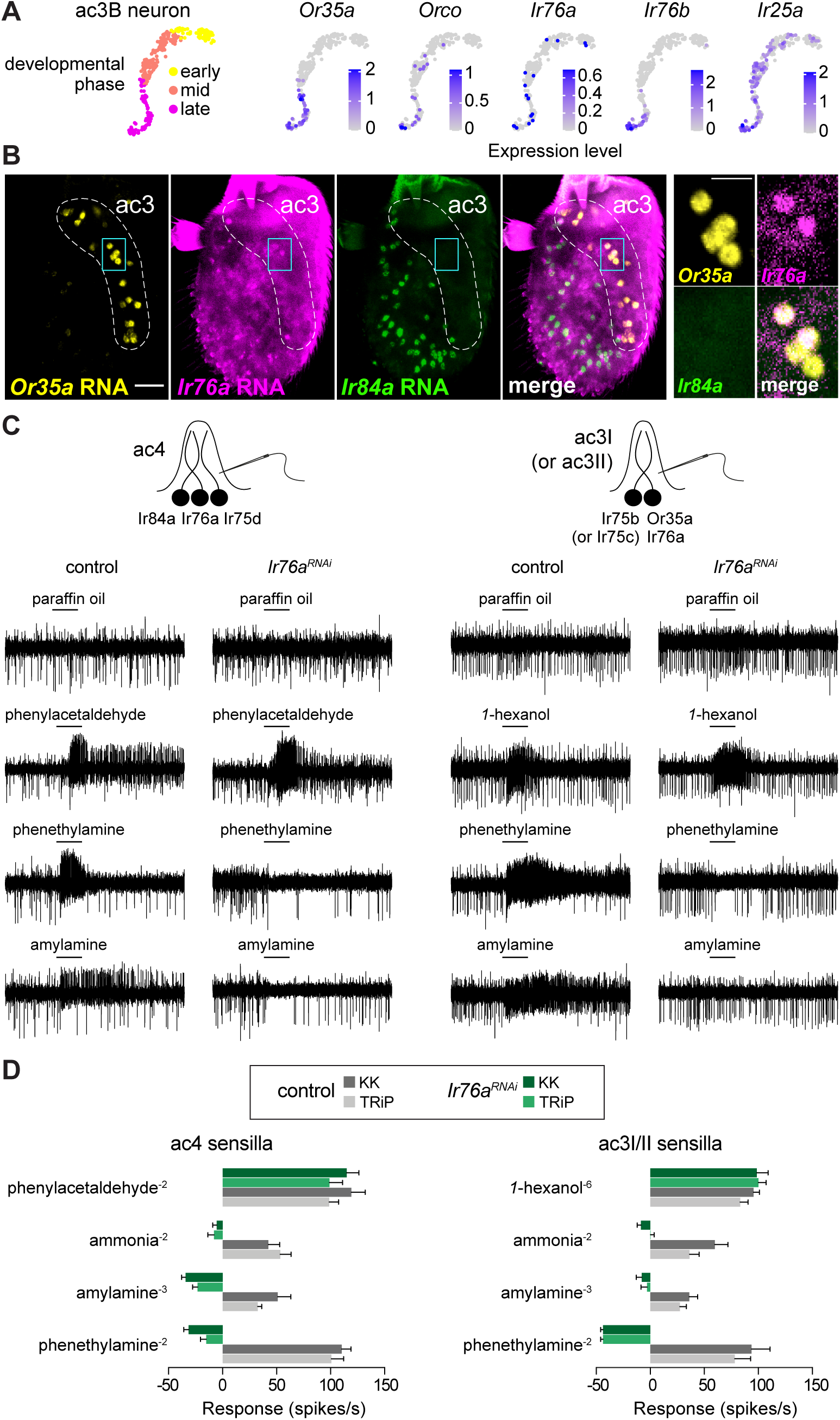
A hybrid Or/Ir OSN population. (A) Top: UMAPs of the ac3B neurons at different development phases extracted from the snRNA-seq atlas (Figure 1A) (Mermet et al., 2025) illustrating the expression patterns of the indicated receptor genes. (B) RNA FISH on a whole-mount antenna of control (*w^1118^*, *n* = 10) animals with probes targeting the indicated transcripts. The ac3 sensilla zone is indicated; distinct from the ac4 zone where Ir84a neurons (and most Ir76a neurons) are located. Scale bar, 25 µm. Right: ac3B neurons co-expressing *Or35a* and *Ir76a* (but not paired with ac4 *Ir84a*-expressing neurons) in a single confocal Z-slice. Scale bar, 10 µm. (C) Representative traces of single-sensillum recordings from ac4 and ac3 sensilla in control and *Ir76a^RNAi^* flies (TRiP lines) illustrating neuronal responses to the indicated odors (0.5 s stimulation time, black bars). (D) Electrophysiological responses to the indicated ligands in ac4 and ac3 sensilla from antennae of two independent lines of control and *Ir76a^RNAi^* animals. Solvent-corrected responses (mean ± SEM) of the combined activities of all neurons in the sensilla are shown (see Data S1 for spike counts, sample sizes and statistical analyses).

The expression of *Ir76a* in ac3B was intriguing because while most odor responses of the broadly-tuned ac3B neuron depend upon Ors (Silbering et al., 2011; Yao et al., 2005), responses to amines – notably phenethylamine and amylamine – require instead the Ir co-receptors Ir25a and Ir76b (Vulpe and Menuz, 2021), which are also expressed in these cells (Figure 3A) (Task et al., 2022). As these amines are amongst the best agonists of ac4 Ir76a neurons (Silbering et al., 2011), we hypothesized that Ir76a is the tuning receptor mediating amine responses in ac3B neurons. We tested this possibility through single-sensillum electrophysiological analyses of *Ir76a^RNAi^* flies (Figure 3C-D). Using two independent transgenic RNAi lines, we first verified the efficiency of *Ir76a^RNAi^* in ac4 sensilla, observing complete loss of responses to amine ligands of Ir76a neurons, while responses of the co-housed Ir84a neurons to phenylacetaldehyde were unchanged (Figure 3C-D). In ac3B neurons, amine responses were similarly abolished by *Ir76a^RNAi^*, while responses to the Or35a/Orco-dependent ligand *1*-hexanol were unaffected (Figure 3C-D).

These results indicate that the ac3B neuron is, to our knowledge, the first unambiguous example of an OSN expressing functionally relevant combinations of tuning and co-receptors of both Or and Ir families. Interestingly, recent snRNA-seq and RNA FISH in the mosquito *Aedes aegypti* identified a few OSN populations in the antenna and maxillary palp expressing putatively complete sets of both Or and Ir complexes (Adavi et al., 2024; Herre et al., 2022), indicating that similar “hybrid” neuron types might exist in other species.

### A new integrated dataset of the developmental, anatomical and functional properties of the *D. melanogaster* olfactory system

Our discoveries of the Or46aB and hybrid Or35a/Ir76a sensory channels both highlighted prior inaccuracies and omissions in the antennal and antennal lobe maps and exemplified the power of using information from disparate sources to extract new insights. We therefore reasoned that it was timely to systematically integrate current data resources on diverse developmental, anatomical and functional properties of the olfactory and hygro/thermosensory systems. Building on a foundational data resource generated nearly a decade ago (Grabe et al., 2016) and from several recent studies on sacculus hygrosensors and thermosensors (Budelli et al., 2019; Enjin et al., 2016; Frank et al., 2017; Gallio et al., 2011; Knecht et al., 2017; Knecht et al., 2016; Marin et al., 2020), we made substantial new additions and corrections regarding receptor expression patterns, neuronal and sensillar annotations. For example, in addition to the definition of ab6 and ab11 described above, we distinguish the classes of antennal intermediate (ai2, ai3) and trichoid (at1, at4) sensilla more clearly, as these have been conflated in the past (e.g., (Couto et al., 2005)). We also update the definition of ac3 sensilla that comprise two subtypes, ac3I and ac3II, housing Ir75b and Ir75c neurons respectively (Prieto-Godino et al., 2017), each together with the Or35a/Ir76a neurons characterized here.

We also collated improved quantitative estimates of neuronal populations favoring numbers from analyses of *in situ* gene expression – including many new quantifications using HCR FISH (Figure S1), other numbers from the literature (e.g., (Mermet et al., 2025)) and from very recent EM connectomic datasets (Dorkenwald et al., 2024; Schlegel et al., 2021; Schlegel et al., 2024) – rather than transgenic reporters as in (Grabe et al., 2016), which do not always faithfully reflect endogenous gene expression. We additionally integrated several developmental properties, such as expression of proneural and other fate determinants, as well as available anatomical information on LNs (Chou et al., 2010) and uniglomerular PNs (Schlegel et al., 2021). Finally, we incorporated comparative datasets of OSN numbers and glomerular size available for several species in the *Drosophila* group (Depetris-Chauvin et al., 2023).

Behavior is of course the *raison d’être* of the olfactory system, and there is a wealth of information on the contributions of many individual olfactory pathways (e.g., (Badel et al., 2016; Semmelhack and Wang, 2009; Wu et al., 2022)). For certain sensory channels, such as those detecting pheromones, several studies provide consistent evidence for their behavioral role(s) (Kurtovic et al., 2007; Taisz et al., 2023). For the majority of pathways, their contribution to odor-evoked behaviors – as assessed by loss-of-function or artificial neuronal activation approaches – are highly context-dependent (Currier and Nagel, 2020), influenced by the experimental assay design (Chin et al., 2018; Tumkaya et al., 2022; Wu et al., 2022), environmental conditions (e.g., air currents (Bell and Wilson, 2016; Matheson et al., 2022; Stupski and van Breugel, 2024)), other simultaneous olfactory and taste inputs (Grabe and Sachse, 2018; Oh et al., 2021; Wilson, 2013) and the internal state of the fly (e.g., starvation (Ko et al., 2015; Lebreton et al., 2015; Root et al., 2011)). Collectively these studies support the idea that many sensory channels function as part of a “combinatorial code” to control behavioral outputs. We have therefore adopted the more general idea of the “sensory scene” within which a particular olfactory pathway might function (Schlegel et al., 2021). This classification is largely defined by the likely ecological source of the odor(s) to which a given OSN responds (Mansourian and Stensmyr, 2015). We caution that such classification is tentative, as some chemicals can be found in many different biological settings.

The full integrated dataset is provided in Table S1; this is also available online (https://shorturl.at/gznii), with the aim that such a dataset can be supplemented with information emerging in future investigations, such as additional molecular markers (McLaughlin et al., 2021; Mermet et al., 2025; Xie et al., 2021), functional properties of individual sensory pathways, and further data from other species of drosophilids (Bontonou et al., 2024). Accompanying this resource, we have created schematics highlighting some key organizational properties of sensory sensilla (Figure 4). We have also generated labeled atlases and movies depicting coronal (anterior-to-posterior) (Figure 5 and Data S2) and transverse (dorsal-to-ventral) (Figure S2 and Data S2) sections through the antennal lobe based on 3D glomerular meshes from a recent EM-based atlas (Bates et al., 2020). Together, these should serve as practical guides during, for example, neurophysiological and anatomical investigations.

**Figure 4.**
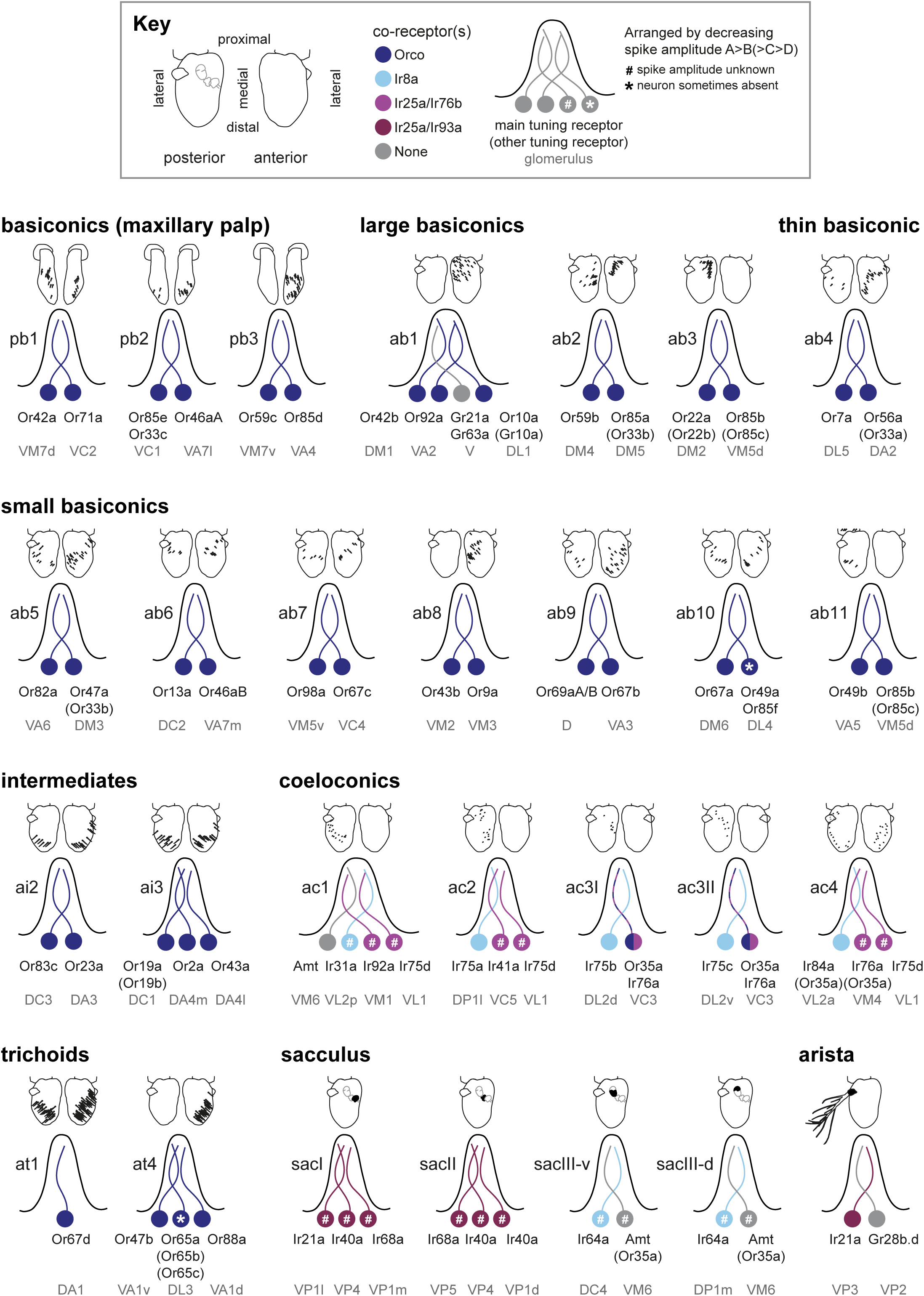
Antennal and maxillary palp sensory sensillum organization. Updated neuronal composition of all sensillar classes in the maxillary palp and antenna, including tuning receptors, co-receptors and the corresponding glomerular targets in the antennal lobe. Tuning receptors shown in parentheses are reported to be expressed in the neuron population but have not yet been shown to contribute to their odor responses; in some cases, these might be non-functional. In ab10 and at4, a specific neuron is sometimes lacking in mature sensilla (asterisks), likely due to promiscuous programmed cell death (Mermet et al., 2025; Nava Gonzales et al., 2021). The approximate distribution of olfactory sensilla within the sensory organs (shown above each sensillum) is adapted from (Grabe et al., 2016) except for ab3 and ab11, which were mapped using image data from (Takagi et al., 2024), and ac3I and ac3II, which were mapped using data from (Mika et al., 2021). While the overall distribution is stereotyped between antennae, there is variation in the individual position of sensilla. The anterior/posterior distribution of large basiconic sensilla does not fully agree with an earlier mapping (de Bruyne et al., 2001), which might reflect differences in definition of the anterior and posterior surfaces between studies.

**Figure 5.**
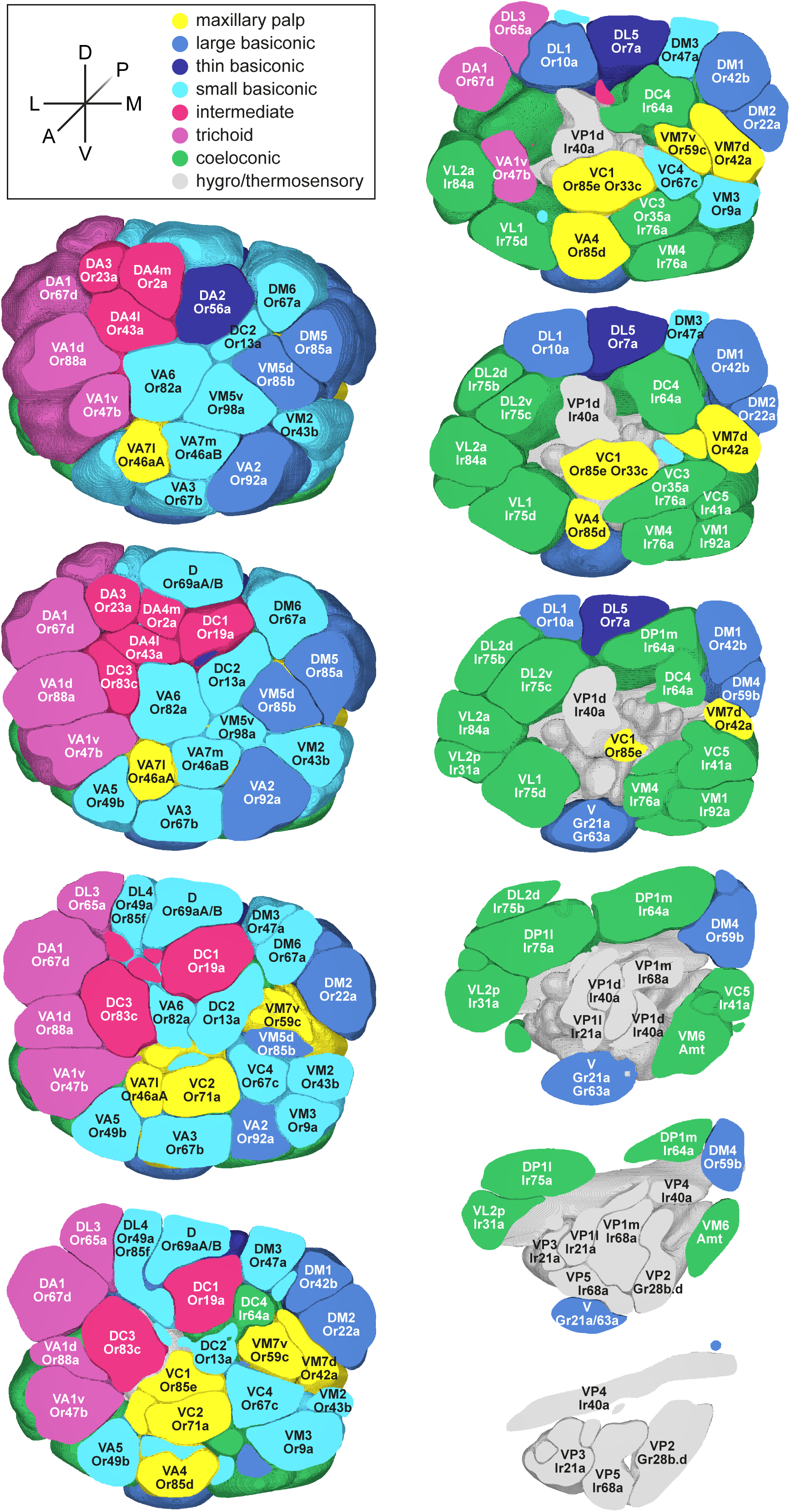
Antennal lobe atlas. Coronal sections through an updated antennal lobe atlas adapted from glomerular meshes based on the female adult fly brain (FAFB) EM dataset (Bates et al., 2020) (see Methods). Anterior is top-left and posterior is bottom-right. The atlas contains updated tuning receptor and glomerular names (Schlegel et al., 2021), and glomeruli are color coded by sensillar class. Glomeruli innervated by OSNs from sacculus chamber III are colored green, as they are most similar to coeloconic neurons. For compactness, only the main known tuning receptor is indicated. For an alternative set of transverse sections along the dorsal-ventral axis, see Figure S2. See Data S2 for an interactive and modifiable version and associated files as well as finer-grained coronal and transverse movies of sections through the antennal lobe.

### Illustration of insights from the integrated dataset

While the information compiled above should serve as a useful reference source during study of specific sensory pathways, we describe in this section a few examples of insights that can be gleaned from global analyses using these updated data.

*Relationship between OSN precursor identity and OSN morphology:* unlike the odor response profile, OSN spike amplitude is not defined by the tuning receptor (Hallem et al., 2004) but rather reflects the morphology of the corresponding OSN. OSNs with greater dendritic surface area, typically due to extensive branching of the sensory cilia endings, have larger spike amplitudes (Nava Gonzales et al., 2021; Shanbhag et al., 1999, 2000). Essentially all sensilla house neurons of distinct, stereotyped spike amplitudes, implying a hard-wired genetic control of neuronal morphology. We asked whether these differences reflect the corresponding neuronal precursor identity. By examining sensilla with two OSNs, we found that the neurons with larger spike amplitudes (A neurons) and those with smaller spike amplitudes (B neurons) were derived from a similar proportion of Nab and Nba precursors (Figure 6A, Table S1). Similarly, in 3-OSN sensilla the A neuron was derived either from Nab (at4, ac2, ac4) or Nba (ai3), and in 4-OSN sensilla the A neuron was derived from either Nba (ab1) or Nbb (ac1). These observations indicate the OSN sensillar morphology is not simply derived from the developmental pathway characteristic of different OSN precursors such as the Notch status after asymmetric cell division (Endo et al., 2007; Endo et al., 2011). Extraction of transcripts enriched in large or small spiking neurons from snRNA-seq datasets (Li et al., 2020; McLaughlin et al., 2021; Mermet et al., 2025) might reveal candidate molecules underlying differences in cilia morphology, an outstanding question in sensory biology in insects and other animals (Maurya, 2022).

**Figure 6.**
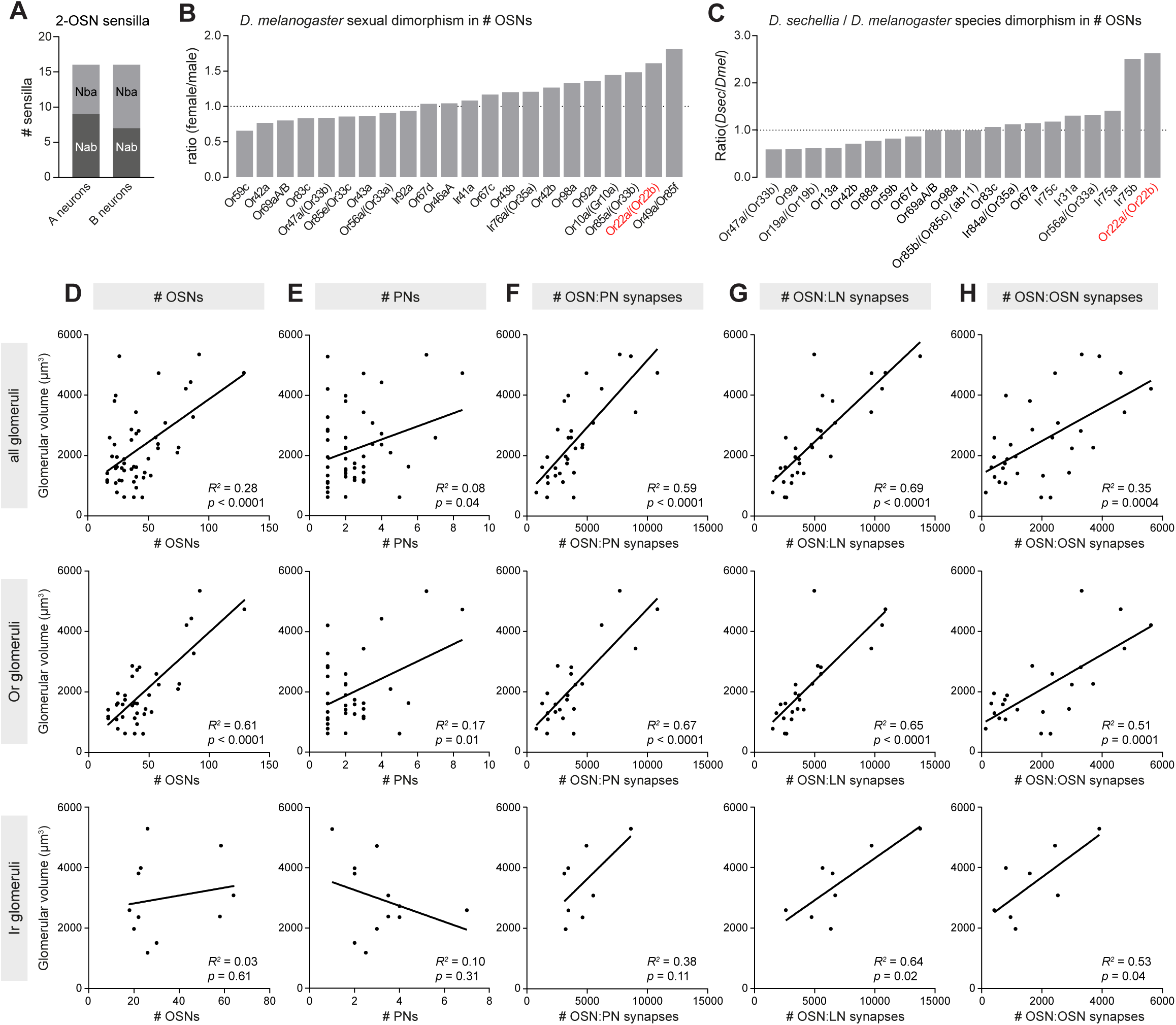
Organizational insights obtained from the resource table. (A) Stacked bar plot of the identity of OSN precursor type (Nab or Nba; Naa and Nbb are absent due to developmental programmed cell death) in large-spike amplitude A and small-spike amplitude B neurons in sensilla with two OSNs. (B-C) Bar plots of the ratio of OSN numbers in female and male *D. melanogaster* and female *D. sechellia* and *D. melanogaster* (C), revealing that the Or22a/(Or22b) population exhibits both sexual and species dimorphism. Note that only OSN populations for which direct experimental data are available (see Table S1) are plotted; however, similar ratios can be inferred for the paired neurons within a given sensillum (e.g., Or85b/(Or85c) neurons in ab3 (Takagi et al., 2024)). (D) Correlation of glomerular volume and OSN numbers for all glomeruli (top), Or glomeruli (middle) and Ir glomeruli including the VC3 Or35a/Ir76a glomerulus (Mermet et al., 2025)(bottom). Note that OSN numbers per glomerulus were used; for nearly all populations this number represents twice the number of OSNs per antenna because most OSNs project bilaterally. There are two exceptions (Ir75d and Gr21a/Gr63a OSNs), which project only unilaterally; here the numbers of neurons per glomerulus are equivalent to those in the antenna. (E) Correlation of glomerular volume and PN numbers for all glomeruli (top), Or glomeruli (middle) and Ir glomeruli (bottom). (F-H) Correlation of glomerular volume and numbers of OSN:PN synapses (F), OSN:LN synapses (G) and OSN:OSN synapses (H) for all glomeruli (top), Or glomeruli (middle) and Ir glomeruli (bottom). For all plots in (D-H), data are from Table S1; coefficients of determination (*R^2^*) and *p* values are indicated on each plot.

*Sexual dimorphisms and species differences in OSN numbers*: many insects have sex-specific olfactory pathways, most famously in moths that possess male-specific populations detecting female pheromones (Nakagawa et al., 2005). By contrast, in *D. melanogaster* sexual dimorphisms in the size of OSN populations appear to be limited. With our revised set of neuron numbers (Table S1), we re-visited this issue by plotting the female:male ratio of OSN numbers, where data are available. While we confirmed that sexual dimorphisms are modest, we noted that sensilla with the greatest over-representation in females are ab10 (implied by greater numbers of Or49a/Or85f neurons) and ab3 (implied by greater numbers of Or22a/(Or22b) neurons) (Figure 6B). Importantly, the latter example was previously overlooked due to underestimation of ab3 numbers quantified using an *Or22a*-Gal4 transgenic reporter (Grabe et al., 2016). The sexual dimorphism in ab3 numbers is noteworthy because these neurons also display interspecific variation in number, notably representing the greatest difference of all Or neuron types between *D. melanogaster* and the ecological specialist *D. sechellia* (Auer et al., 2021), which has 2-3-fold more ab3 OSNs (Auer et al., 2020; Dekker et al., 2006; Takagi et al., 2024) (Figure 6C). We recently provided evidence that increased OSN population size in *D. sechellia* enhances olfactory behavior not by increasing sensitivity of partner PNs, but rather by influencing their adaptation properties to repetitive or prolonged stimuli (Takagi et al., 2024). This invites the question of whether the dynamics of odor processing in PNs receiving input from ab3 and ab10 neurons are sexually dimorphic in *D. melanogaster* due to the differences in OSN number.

Shared sexually dimorphic and interspecific differences in OSN population size are not observed for other populations. For example, while ab10 Or49a/Or85f neurons are over-represented in females, there is no species difference in ab10 (as inferred from Or67a OSN numbers) between *D. melanogaster* and *D. sechellia* (Figure 6B-C). Reciprocally, while the ac3I Ir75b neuron population is greatly expanded in *D. sechellia* compared to *D. melanogaster* (Figure 6C), it is of a similar size in males and females in both species (Prieto-Godino et al., 2017; Takagi et al., 2024).

*Relationship of glomerular size with neuron and synapse numbers*: previous studies suggested a shallow, but significant correlation between the number of OSNs and the size of the corresponding glomerulus (Grabe et al., 2016). We re-analyzed this relationship, both for all glomeruli where data is available, and those receiving input from Or and Ir OSNs separately (Figure 6D). While we confirmed a statistically significant correlation overall, we found that this is driven by a strong relationship with Or glomeruli, as Ir OSN number and glomerular size are uncorrelated (Figure 6D). These observations indicate that Ir glomerular size must be dictated by other properties.

Using the more extensive dataset from the FlyWire connectome (Dorkenwald et al., 2024; Schlegel et al., 2024), we therefore examined correlations between glomerular size and PN number, but there was no evidence of a strong relationship, globally or within either olfactory subsystem (Figure 6E). However, comparison of glomerular size with the number of synapses that individual classes of OSNs make with PNs, LNs, and other OSNs in the hemibrain connectome (Schlegel et al., 2021) revealed positive correlations in all cases, although this was only a trend for Ir glomeruli for OSN:PN synapses, potentially because of limited sample size (Figure 6F-H). These observations indicate that the densities of OSN:PN, OSN:LN and OSN:OSN synapses are relatively consistent across glomeruli regardless of the number of input or output neurons. The determinant of Ir glomerular size differences remains an interesting open question, which might be answered by future analysis of other microarchitectural features revealed by the connectome.

## Concluding remarks

Through identification of new olfactory sensory channels in *D. melanogaster*, we have “completed” our understanding of the basic molecular organization of this sensory system, notwithstanding structural and functional heterogeneity that undoubtedly exists within at least some sensory pathways. Using this finding as a stimulus to create an updated, integrated data resource of much of the enormous body of knowledge of the construction and function of this species’ olfactory (as well as hygrosensory and thermosensory) systems, we believe this work should facilitate and inspire the coming years of research in the field.

## Methods

### RNA FISH

HCR RNA FISH was performed on a control *peb-Gal4* genotype (RRID:BDSC_80570) (Figures 1-2, Figure S1) or *w^1118^* (Figure 3) using female flies, as described (Mermet et al., 2025). All probes were produced by Molecular Instruments (Table S2). Images from antennae and maxillary palps were acquired with confocal microscopes (Zeiss LSM710 or Zeiss LSM880 systems) using a 40× (or 63× for the palp) oil immersion objective and processed using Fiji software (Schindelin et al., 2012).

### Electrophysiology

GFP-guided single sensillum electrophysiological recordings were performed on 2-day old females using glass electrodes filled with sensillum recording solution, essentially as described (Vulpe et al., 2021). For ab6 sensilla we used *Or13a-Gal4/UAS-mCD8::GFP* (parental stocks RRID:BDSC_23886 and RRID: BDSC_5130); for ab11 sensilla we used *Or49b-Gal4*/*UAS-mCD8::GFP* (parental stocks RRID:BDSC_24614 and RRID:BDSC_5130). A Prior Scientific Lumen 200 Illuminator was used as the excitation light source. The sample was visualized using a BX51WI Olympus microscope with a 1.6× magnification changer, a 50× objective and a Semrock GFP-4050B-OMF filter cube.

For *Ir76a* loss-of-function analysis in ac3 and ac4, we crossed the *P{Act5C-GAL4}25FO1* driver (RRID:BDSC_4414) to the following *Ir76a^RNAi^* or RNAi control transgenic lines: *UAS-Ir76a^RNAi^*(KK) (VDRC_101590), *UAS-Ir76a^RNAi^* (TRiP) (RRID:BDSC_34678), RNAi control (KK) (VDRC_60100), RNAi control (TRiP) (RRID:BDSC_36303) (see Data S1 for final genotypes). ac3 and ac4 sensilla were identified based upon their stereotyped location on the antenna (Benton et al., 2009) and their responses to diagnostic odors (Silbering et al., 2011).

Odorants (Table S3) were diluted (v/v) in paraffin oil (or water for ammonia), as indicated in the figure plots. Odor cartridges were prepared by applying 50 µl odorant solution onto a Whatman 13 mm assay disc, which was inserted into a Pasteur pipette closed with a 1 ml pipette tip. Fly preps were placed in a 2 l/min air flow directed by a glass air tube. Odor stimuli were injected into the air flow for 0.5 s at 0.5 l/min. The odor response was calculated from the difference in OSN spike frequency (or summed frequencies of all OSNs for ac sensilla) in response to a 0.5 s odor puff compared to a 0.5 s solvent puff, as described (Vulpe et al., 2021).

### Terminology

There is some inconsistency in the literature regarding the use of certain terms, which we aim to clarify here.

First, “Olfactory Receptor Neuron” (ORN) and “Olfactory Sensory Neuron” (OSN) terms have been used interchangeably. We have favored the latter, as the terminology “sensory” describes more generally the function of these neuron populations, rather than linking them to a molecular entity (“receptor”). Moreover, this general terminology better encompasses the diversity of sensory neuron types, which can express Ors, Irs or Grs.

Second, the use of the terms “tuning receptor” and “co-receptor” are generally well-accepted, though not equally applicable in every neuron. “Tuning receptor” refers to the subunit defining stimulus-specificity of a sensory receptor complex, and likely directly binds and/or is conformationally modified by the stimulus. Some neurons house multiple potential tuning receptors; the best-characterized case is the maxillary palp pb2 neuron expressing two functional receptors, Or85e and Or33c (Goldman et al., 2005). Several other cases of tuning receptor co-expression have been described, but only one receptor is functional (e.g., the ab4 neuron expressing Or56a and Or33a, where only the former receptor appears to contribute to neuronal specificity (Stensmyr et al., 2012)). In this study we indicate such potentially non-functional receptors in parentheses. “Co-receptors” are obligatory subunits necessary for olfactory receptor trafficking and function. Due to their broad expression across multiple classes of neurons, they are assumed not to contribute to the sensory specificity of a particular neuron type and likely do not bind ligands; while this is clearest for the Or co-receptor Orco, several Ir co-receptors exhibit narrower expression patterns in sets of neurons that respond to particular functional classes of stimuli (e.g., Ir76b in amine-sensing neurons; Ir93a in hygro/thermosensory neurons), and it cannot be excluded that such proteins have a more direct role in stimulus recognition. Many co-receptors are expressed in neurons where there is no corresponding tuning receptor (Task et al., 2022), but there is so far little evidence for their roles in such neurons (see also (Mermet et al., 2025)). Finally, tuning and co-receptor identity is ambiguous or irrelevant in certain neurons. For example, in aristal Gr28b.d neurons, this Gr appears to function alone (Mishra et al., 2018; Ni et al., 2013). In ab1C CO_2_-sensing neurons, both Gr21a and Gr63a are, at least in *Xenopus* oocytes, partially sufficient for conferring sensory responses, although less effectively than these receptors together (Ziemba et al., 2023), and both are required for *in vivo* reconstitution of CO_2_ sensitivity in heterologous neurons (Jones et al., 2007; Kwon et al., 2007).

Third, for sensillum nomenclature, we note the literature contains several discrepancies in the descriptions of the neuronal composition of ab6 and ai1 sensilla. The first characterization of ab6 was through electrophysiological recordings, which demonstrated the presence of two neurons: one responded to various alcohols (notably *1*-octen-*3*-ol) and the other to *4*-methylphenol (de Bruyne et al., 2001). Subsequent functional studies matched the response profile of Or49b receptors to ab6B neurons (Hallem et al., 2004). Further molecular and histological studies tentatively suggested Or49b is housed in the ab6 sensillum with Or85b and/or Or98b neurons (Couto et al., 2005). However, a later survey proposed that Or49b and Or13a neurons are paired in this sensillum, due to the close similarity of Or13a and ab6A response profiles (Galizia et al., 2010). This proposition was re-quoted in subsequent papers (e.g., (Auer et al., 2020; Grabe et al., 2016; Prieto-Godino et al., 2020)). Concurrently, targeted recording of sensilla housing Or13a neurons (through expression of GFP under the control of *Or13a*-Gal4) lead to its designation as the sole neuron housed in so-called ai1 sensilla, distinct from “ab6” sensilla housing Or49b neurons (Lin and Potter, 2015). However, the length of the putative ai1 sensillum resembles more closely small basiconic sensilla than other ai sensilla (Lin and Potter, 2015). Moreover, our re-analysis of electrophysiological traces from that study (Lin and Potter, 2015) revealed the presence of two spike amplitudes in at least some sensilla (data not shown), and our new recordings (Figure 2B) unambiguously demonstrate the presence of a second neuron in this sensillum, which we have shown expresses Or46aB.

Recently, we demonstrated that Or49b-expressing neurons are paired with those expressing Or85b/(Or85c), and we described these as ab6 sensilla based on their expression of Or49b (Takagi et al., 2024). This receptor pairing might have been overlooked in previous studies because the majority of Or85b/(Or85c) neurons are housed in ab3, paired with Or22a/(Or22b) neurons (Takagi et al., 2024). In the current study, we have determined that there are two sensilla populations that could potentially be named ab6: those housing Or49b and Or85b/(Or85c) neurons and those with Or13a and Or46aB neurons. We propose to give precedent to the original electrophysiological analysis (de Bruyne et al., 2001) by designating the ab6 sensillum as that housing Or13a and Or46aB neurons. The sensillum housing Or49b and Or85b/(Or85c) neurons therefore represents a new type of sensillum, which we name ab11. Finally, we note that one report described “ab11” and “ab12” sensilla, each housing three OSNs, one of which responds to the insect repellent citronellal (Kwon et al., 2010). The molecular identity of these sensilla is unclear, and they have not been described in any subsequent studies. Given the apparent completeness of the antennal lobe map with our discovery of Or46aB neurons, we suggest the sensilla classes described in that study represent variants of other basiconic classes (e.g., a three-OSN “abX” from (Nava Gonzales et al., 2021)), rather than new classes.

### Data resources and analysis

The snRNA-seq data and analysis methods are described in (Mermet et al., 2025); gene expression levels shown in the UMAPs are residuals from a regularized negative binomial regression, and have arbitrary units. The antennal lobe confocal images are from (Endo et al., 2007). The antennal lobe atlas used glomerular meshes previously generated by EM analysis of the antennal lobe (Bates et al., 2020), incorporating updated glomerular naming (Schlegel et al., 2021). Antennal lobe images were generated using the open-source software 3D Slicer (Fedorov et al., 2012) (see Data S2). Statistical analyses and plots were generated in RStudio with Seurat (v4.3.0.1) and GraphPad Prism 10.3.1. All other main sources of data are referenced directly in Table S1.

## Acknowledgements

We are grateful to Roman Arguello for sharing bulk-RNA datasets, to Chihiro Hama for sharing antennal lobe image data, and Veit Grabe and Silke Sachse for sharing anatomical data and discussions. We thank Tom Auer, Jamie Jeanne, Chris Potter, Lucia Prieto-Godino, Marcus Stensmyr, Chih-Ying Su, and members of the Benton and Menuz laboratories for feedback. Research was supported by NIH awards R35GM133209 and R21DC021267 to K.M. Research in R.B.’s laboratory is supported by the University of Lausanne, an ERC Advanced Grant (833548) and the Swiss National Science Foundation (310030 219185).

## Author contributions

R.B. conceived and supervised the project and collated and analyzed most data for Table S1. J.M. identified and characterized the Or46aA/B and Or35a/Ir76a cell types through snRNA-seq analysis and RNA FISH and contributed other OSN population quantifications. A.J. performed and analyzed electrophysiological experiments. K.E. provided data from SOP lineage labelling experiments. S.C. performed and quantified RNA FISH experiments. K.M. conceived and supervised the project, contributed to data collation in Table S1, analyzed bulk RNA-seq and generated the antennal lobe atlas files. R.B. and K.M. prepared the figures, with contributions from J.M. and A.J. The manuscript was drafted by R.B. with input from K.M. and J.M. All authors approved the final manuscript.

## Supplementary Tables and Figures

**Table S1.** Maxillary palp and antennal neuron cell types and circuitry.

**Table S2.**
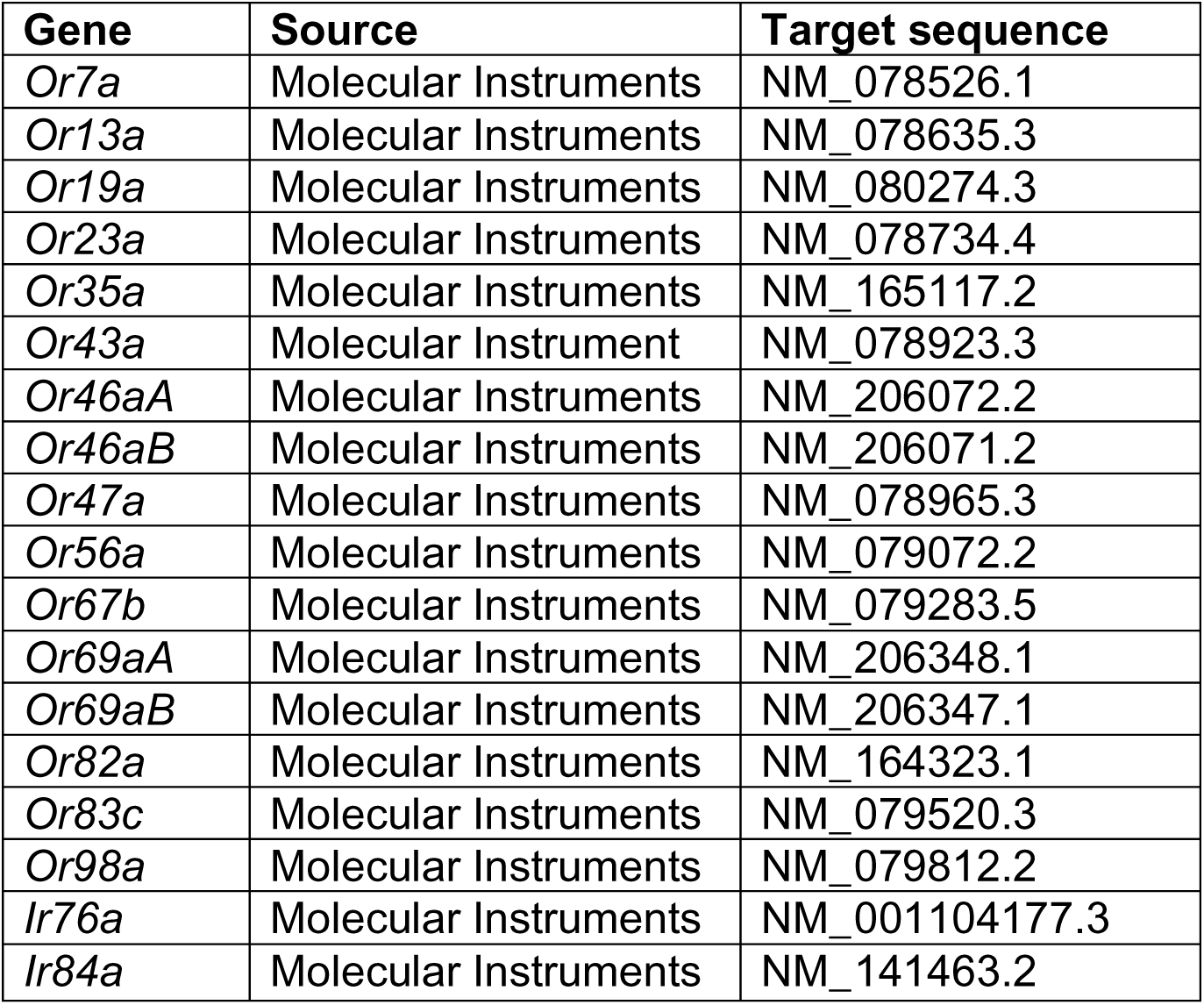
RNA FISH probes.

**Table S3.**
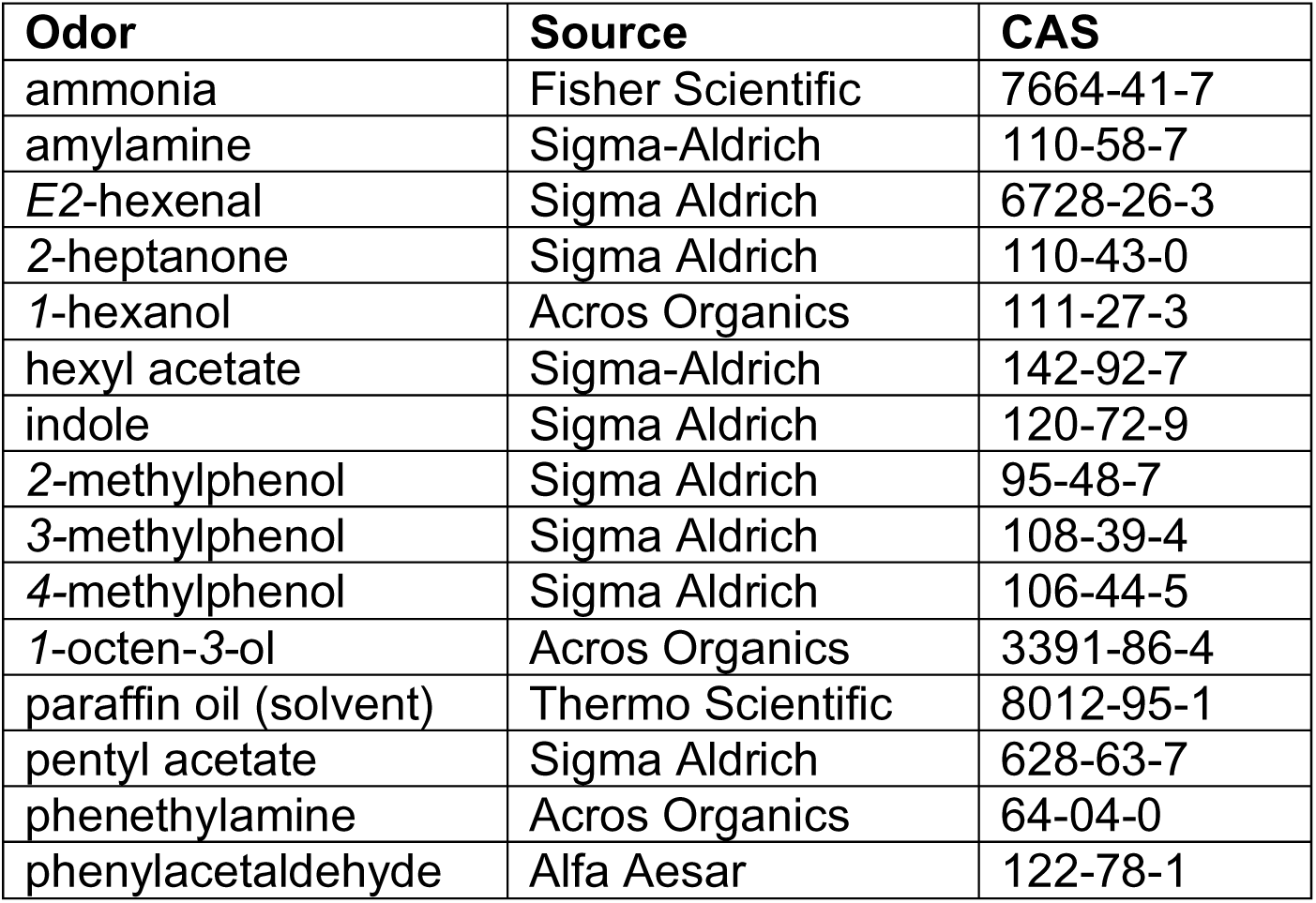
Odors.

| Odor | Source | CAS |
| --- | --- | --- |
| ammonia | Fisher Scientific | 7664-41-7 |
| amylamine | Sigma-Aldrich | 110-58-7 |
| *E2*-hexenal | Sigma Aldrich | 6728-26-3 |
| 2-heptanone | Sigma Aldrich | 110-43-0 |
| 1-hexanol | Acros Organics | 111-27-3 |
| hexyl acetate | Sigma-Aldrich | 142-92-7 |
| indole | Sigma Aldrich | 120-72-9 |
| 2-methylphenol | Sigma Aldrich | 95-48-7 |
| 3-methylphenol | Sigma Aldrich | 108-39-4 |
| 4-methylphenol | Sigma Aldrich | 106-44-5 |
| 1-octen-3-ol | Acros Organics | 3391-86-4 |
| paraffin oil (solvent) | Thermo Scientific | 8012-95-1 |
| pentyl acetate | Sigma Aldrich | 628-63-7 |
| phenethylamine | Acros Organics | 64-04-0 |
| phenylacetaldehyde | Alfa Aesar | 122-78-1 |

**Figure S1.**
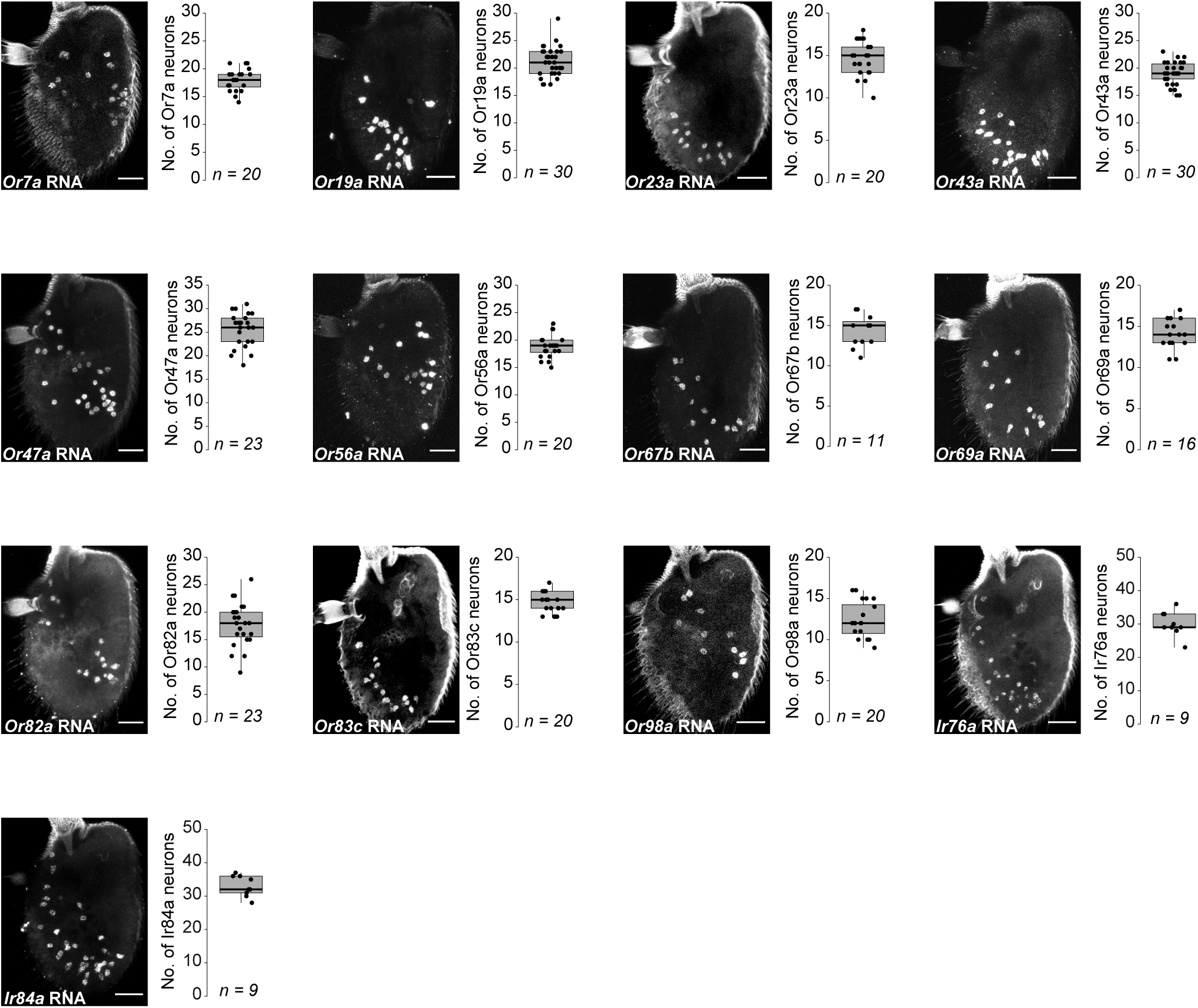
Quantification of OSN populations by HCR RNA FISH. Representative images of HCR RNA FISH on whole-mount antennae (control genotype *peb-Gal4*) using the indicated gene probes, and quantifications of OSN population size. These data were used to complement information in Table S1. For Or69aA/B neurons, the image shown is with an *Or69aA* probe, but the quantifications are pooled from images using either *Or69aA* or *Or69aB* probes. Scale bars, 25 μm.

**Figure S2.**
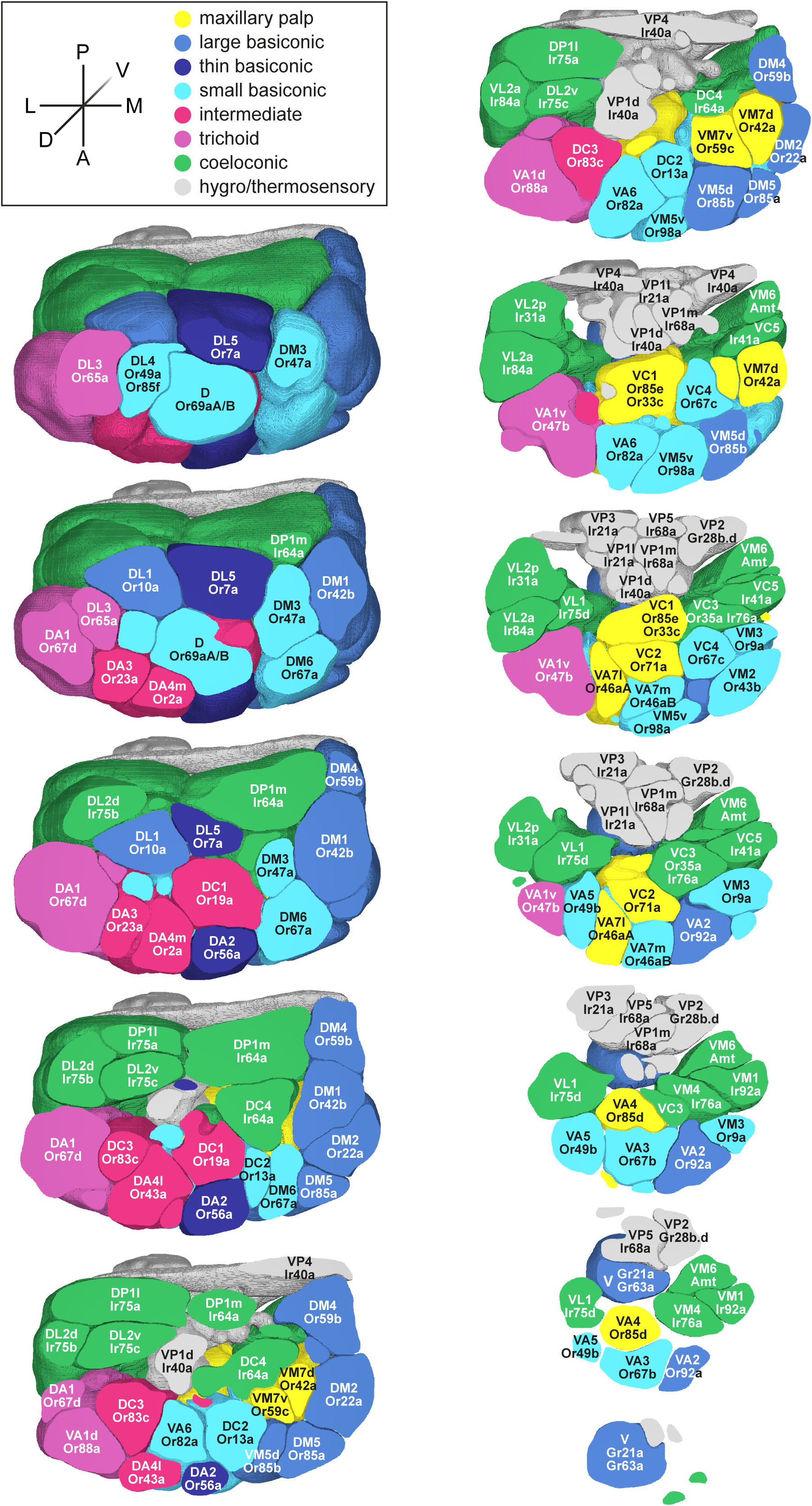
Antennal lobe atlas. Transverse sections, along the dorsal-ventral axis, of the antennal lobe atlas shown in Figure 5. Such views are more typical of those obtained during *in vivo* calcium imaging experiments.

## Supplementary Datasets

**Data S1. Odor-evoked neuronal responses.**

**Data S2. Antennal lobe atlas data.**

The DataS2.seg.vtm file and associated folder contain segmentations of the antennal lobe created from the glomerular mesh models generated previously from the female adult fly brain (FAFB) connectome (Bates et al., 2020). When opened in the open-source software 3D Slicer (Fedorov et al., 2012) and viewed in the Segmentation module, the antennal lobe can be viewed in 3D with each glomerulus represented as an individual segmentation labeled with its name, and associated receptor(s) and sensillum. The antennal lobe can be rotated for viewing from different angles, and interactive coloring and visibility control of individual glomeruli are supported. A binary labelmap was created from DataS2.seg.vtm and used to generate the DataS2-label.nrrd volume file and the color lookup file DataS2-label_ColorTable.ctbl. When opened together in 3D Slicer and viewed in the Volume module, the colors from DataS2-label_ColorTable.ctbl can be associated with the glomeruli in the DataS2-label.nrrd volume. When viewed in the Volume Rendering module, the 3D volume can be visualized and slices taken through the volume (anterior-to-posterior, lateral-to-medial or dorsal-to-ventral axes) using the ROI feature. Data S2 also contains two movies of coronal (anterior-to-posterior) and transverse (dorsal-to-ventral) slices through the antennal lobe colored as in Figure 5 and DataS2.seg.vtm.

